# Evolution of transcription factor-containing superfamilies in Eukaryotes

**DOI:** 10.1101/2024.09.24.614687

**Authors:** Akshara Dubey, Ganesh Muthu, Aswin Sai Narain Seshasayee

## Abstract

Regulation of gene expression helps determine various phenotypes in most cellular life forms. It is orchestrated at different levels and at the point of transcription initiation by transcription factors (TFs). TFs bind to DNA through domains that are evolutionarily related, by shared membership of the same superfamilies (TF-SFs), to those found in other nucleic acid binding and protein-binding functions (nTFs for non-TFs). Here we ask how TF DNA binding sequence families in eukaryotes have evolved in relation to their nTF relatives. TF numbers scale by power law with the total number of protein-coding genes differently in different clades, with fungi usually showing sub-linear powers whereas chordates show super-linear scaling. The LECA probably encoded a complex regulatory machinery with both TFs and nTFs, but with an excess of nTFs when compared to the relative distribution of TFs and nTFs in extant organisms. Losses drive the evolution of TFs and nTFs, with the possible exception of TFs in Animalia for some tree topologies. TFs are highly dynamic in evolution, showing higher gain and loss rates than nTFs though both are conserved to similar extents. Gains of TFs and nTFs are driven by the appearance of a large number of new sequence clusters in a small number of nodes, which determine the presence of as many as a third of extant TFs and nTFs as well as the relative presence of TFs and nTFs. Whereas nodes showing explosion of TF numbers belong to multicellular clades, those for nTFs lie among the fungi and the protists.

## Introduction

Regulation of gene expression contributes to phenotype establishment and to efficient and economic operation of most, if not all, forms of cellular life. It helps determine when genes are expressed and at what levels. Differences in the expression of orthologous genes between closely-related organisms can cause consequential trait divergence. Examples include gene expression differences among primates (1) contributing, for example, to differences in skeletal tissue development between chimpanzees and humans (2). In bacteria, changes in gene regulation can cause stark differences in host specificity between members of the same species (3). In multicellular organisms, large differences in the structure and function of genetically identical tissue types arise from differences in gene expression.

Gene expression is regulated at several levels, from regulation of transcription initiation, elongation and termination, through extensive post-transcriptional control of RNA stability and translation, to modulation of protein activity and turnover. Transcription factors (TFs) are DNA-binding proteins that specifically recognise and bind to certain short DNA sequence (or structural) motifs and, on binding, activate or repress transcription of a proximal gene. Across domains of life, TFs differ from other modulators of transcription, such as those that structure the chromatin or the nucleoid, in usually displaying stronger specificity in binding to particular DNA sequence motifs (4). TFs typically belong to a small set of sequence families (5). This allows us to computationally predict and catalogue the vast majority of TFs encoded in any sequenced genome. For example, most TFs in bacteria belong to a subset of sequence families representing the Helix-Turn-Helix (HTH) domain (6). Eukaryotes have evolved and expanded a distinct set of TF sequence families. These include some subtypes of the homeodomain, which is related to the prokaryotic HTH, and some varieties of the zinc finger family, which is similar to the zinc beta ribbon domain found in a small number of nucleic acid binding proteins in prokaryotes (7). Taken together, sequence family-based surveys of TFs across sequenced genomes indicate that these proteins are encoded in large numbers (∼300 in *Escherichia coli* and ∼1,500 in *Homo sapiens*) in most cellular organisms. The number of TFs encoded by a genome scales positively with the total number of protein coding genes (8) (quadratically in bacteria and more linearly in eukaryotes).

Despite the abundance of TF-encoding genes and their roles in phenotype determination, TFs are not fundamental to all cellular life. Early comparative genomic approaches for defining a minimal genome for cellular life showed that TFs are not conserved universally (9, 10). Later research showed that TFs in bacteria are subject to widespread horizontal gene transfer (11), arguing against earlier work favouring a predominant role for gene duplication (12) in their expansion. At some phylogenetic resolutions, TFs, on average, are less conserved than non-TFs (13). However, TFs may not necessarily be much less conserved than their own targets (14). Among bacteria, the evolution of endosymbiont genomes is driven by gene loss and TFs are lost more frequently than many other types of genes; in fact, many such genomes do not code for any TF (15). Similarly, among eukaryotes, TFs are relatively few in number in parasitic organisms such as *Plasmodium* (16). Thus, the need for TFs appears to arise as organisms evolve and maintain ‘complex’ and diverse lifestyles manifested in genomes encoding large numbers and varieties of genes and gene functions.

How and when did TFs evolve and their repertoires expand? Much work has gone into the evolution of TF specificity and function in various eukaryotic clades (17). For example, recent work suggests that the binding specificity of orthologous TFs can diverge greatly between humans and flies, contrary to claims from older literature (18). At deeper phylogenetic levels, we have evidence suggesting that certain families of TFs, such as the MYB, were likely present in the last common ancestor of eukaryotes (19). Previous work by Vaquerizas et. al. (2009) analysing the conservation of cross 24 genomes including human TFs across 24 different eukaryotes *Homo sapiens* showed the emergence of different TF families at different stages of evolution of human lineage (20). A more recent work by de Mendoza et al. (2013), incorporating a eukaryotic phylogeny into the analysis, had traced the evolution of TFs across eukaryotes up to the last common ancestor and showed the presence of multiple TF families at the common ancestor of eukaryotes and richer TF repertoires in multicellular clades (19). A study of metazoan TFs by Schmitz et. al (2016) highlighted the role of single-gene and while-genome duplications to TF expansions (21).

Despite these studies, the question of the emergence and expansion of TF function across eukaryotes, *in comparison with related families with non-TF functions*, remains less well explored. This relationship between TFs and their non-TF relatives is worth exploring for the following reason. Based on studies of CRP, a global TF in *E. coli* that binds to hundreds of sites, including those with little potential for proximal control of transcription, and the often vague boundary separating TFs from chromosome-shaping nucleoid-associated proteins in bacteria, Busby and Visweshwaraiah argued that sequence-specific TFs evolved from non-specific chromatin/nucleoid-associated proteins involved in more fundamental roles in chromosome compaction (4). Sequence specificity and other TF-associated functions, such as signal sensing and interfaces to interact with the RNA polymerase, could have evolved on top of non-specific chromatin-binding protein sequences. For example, some sequence families, such as that for HU/ IHF, include both non-specific and highly specific DNA binding proteins. There is also evidence that a single amino acid change can convert a promiscuous DNA binding protein into one that displays ‘exquisite’ sequence specificity (22, 23). A TF can be coerced to alter its specificity with one or just a few amino acid substitutions(24). These suggest that evolution of sequence specific TFs from non-specific DNA binding proteins and vice-versa are not difficult to achieve.

In this study, we ask how and when the repertoire of TFs and related non-TFs encoded by eukaryotic genomes expanded. In particular, we ask how TF and non-TF counts scale with genome size across eukaryotes; what the complement of TFs and non-TFs in the LECA was; once acquired, how dynamic are TFs and their non-TF relatives; and whether TF and their non-TF relatives evolved differently and at different stages on the eukaryotic phylogeny.

## Methods and Materials

### Data

The list of 1271 eukaryotic organisms was obtained from OrthoDB database version 10.1 (25). Fasta sequences for these organisms were extracted from odb10v1_all_fasta.tab file using an in-house Python script. This file contains the amino acid sequence for the longest isoform of each gene. This file was used to calculate the proteome size, or the number of protein-coding genes encoded in a genome, of the organisms. Classification of organisms into their respective kingdoms and phyla was obtained from the NCBI taxonomy browser (26) by matching taxid against odb10v1_species.tab. The final organism list contains 148 protists (not a monophyletic group, but a convenience classification), 549 fungi, 121 plants and 448 animals. The list of organisms is present in Supplementary dataset 1.

### DBD prediction

The initial list of DNA binding domains (DBDs) families, in the form of PFAMs, was obtained from Vaquerizas et al. (20) and transcriptionfactor.org (website not accessible now)(5). These were curated using a survey of the literature to assemble a set of 275 DBDs. These domains were then classified, based on a literature survey, into TFs and non-TFs (nTF) families. It is to be noted that domains performing regulatory functions via chromatin remodelling are considered as nTFs. Domains that lacked information about their function were classified as “uncategorised”. These PFAMs (27) were grouped into their corresponding superfamilies depending on the information available on the Superfamily database (28), SCOPS (29) classification and Interpro (30) classification. These superfamilies are called TF-SFs for TF-containing superfamilies in this study. Those PFAMs that did not fall into any superfamily were marked as “unclassified”. This resulted in a total of 159 TFs, 104 nTFs and 12 uncategorised families belonging to 34 different superfamilies. 50 families were marked as unclassified. MYB and SANT proteins, the former being a TF and the latter not, both belong to the same large family. This family was further sub-classified into MYB and SANT subfamilies on the basis of Prosite information available in the Interpro database. The corresponding Interpro ID (version 5.36) for all families and superfamilies (Supplementary dataset 2) were also obtained.

**Table 1:** Statistics on dataset used in this study.

| S No. | Dataset | Number of organisms | TF-SF count | TF family count | nTF family count | Total protein count | TF protein count | nTF protein count | Uncategorized protein count | TF domain count | nTF domain count | TF OG count | nTF OG count |
| --- | --- | --- | --- | --- | --- | --- | --- | --- | --- | --- | --- | --- | --- |
| a. | All TF-SFs; all organisms in dataset | 1271 | 32 | 137 | 79 | 884266 | 681942 | 143166 | 59158 | NA | NA | NA | NA |
| b. | TF-SFs representing both TF and nTF families; all organisms in dataset | 1271 | 14 | 70 | 79 | 678613 | 480760 | 143166 | 54687 | NA | NA | NA | NA |
| c. | All TF-SFs; organisms present in 18S rRNA trees | 704 | 31 | 133 | 77 | 442013 | 337909 | 74379 | 29725 | 640723 | 163549 | 28493 | 5693 |
| d. | TF-SFs representing both TF and nTF families; organisms present in 18S rRNA trees | 704 | 13 | 67 | 77 | 338003 | 236333 | 74379 | 27291 | 482662 | 163549 | 21603 | 5693 |
| e. | All TF-SFs; organisms present in euBac-concat tree | 423 | 31 | 127 | 70 | 328631 | 257366 | 51793 | 19472 | 640721 | 163544 | 24829 | 4661 |
| f. | TF-SFs representing both TF and nTF families; organisms present in euBac-concat tree | 423 | 13 | 62 | 70 | 259554 | 189423 | 51793 | 18338 | 482660 | 163544 | 19243 | 4661 |

### TF prediction and classification

Interproscan standalone software version 5.36 was used to predict the domain / sequence family in the protein sequences (31). HMM profiles from PFAM and Superfamily databases were used to predict these domains using the default settings. Each protein was classified into TFs, nTFs and unclassified depending upon the type of families present. Those proteins that contained a TF-SF but no family annotation were marked as uncategorised. Proteins that harboured a TF family along with nTF or unclassified families were labelled as TFs and those with nTFs and unclassified families as nTFs (Supplementary dataset 3).

### TF-SF protein count v/s proteome size analysis

TF and nTF counts were calculated for each of the 1271 eukaryotic organisms using an inhouse python script (Supplementary dataset 3). The corresponding proteome size was calculated by counting the number of proteins for each organism provided in the odb10v1_all_fasta.tab file. Using the NCBI taxonomy information the slope and the intercept was calculated for log_10_ proteome size vs. log_10_ TF or nTF counts plot for the four kingdoms and also for major phyla within these kingdoms. The HuberRegressor() function from the sklearn python module was used for the same.

### Phylogenetic tree constructions

The *18S-archaea* tree was made using 736 eukaryotic 18S rRNA sequences and 1 archaeal 16S rRNA sequence obtained from the SILVA database version 132(32). The set was limited to the 700+ organisms for which 18S rRNA were available in the database; further removal ensured species-level non-redundancy in our dataset. The tree was constructed using the SILVA ACT online web server with GTR as the evolutionary model and GAMMA as the rate model for the likelihoods (33). The tree obtained was then pruned using an inhouse R script by dropping 32 organisms, resulting in 704 eukaryotes and 1 archaea, which was also used to root the tree in iTOL (34).

To test multiple topologies, *euBac-18S* and *euArch-18S* trees were constructed using constraints on the 18S rRNA alignments. Two different phylogenetic topologies were used as constraints. The 2 trees used as the constraint for *euBac-18S* and *euArch-18S* trees respectively were obtained from Derelle et al. (PNAS 2014) (35) and Jewari et al. (Science 2023) (36) and were pruned for organisms overlapping with our list. To make the constrained *euBac-18S* and *euArch-18S* trees, a trimmed 18S rRNA sequence alignment of 704 eukaryotic organisms was obtained from the SILVA ACT online web server. This alignment was then used as input for iqtree version 2.2.2.6 with GTR as the evolutionary model and an ultrafast bootstrap of 1000 and “-g” option to provide the corresponding constraint tree (37). The *euArch-18S* tree was further pruned by dropping *Amphimedon queenslandica*, thereby resulting in 703 organisms. Both the trees were rerooted similar to their respective constraint trees using iTOL. Across the three 18S rRNA trees at least 60-70% of the internal branches have a bootstrap confidence >= 80% with deep nodes (ones close to the root) showing >= 90 % bootstrap confidence (Fig S1 A,B,C).

For the *euBac-concat* tree, an orthologous gene-based approach was employed. 25 eukaryotic genes of bacterial ancestry (euBac) reported by He et al., 2014 (38) were used across 868 eukaryotes and one bacterium, *Rickettsia canadensis,* which was used to root the phylogenetic tree. MSA of each euBac protein set was then performed using MUSCLE v3.8.31(39), and the conserved regions relevant for phylogenetic inference were trimmed using BMGE (40). The trimmed euBac multi-fasta files were then concatenated using a Perl script [https://github.com/nylander/catfasta2phyml]. A gene partition file was used to specify the start and end positions of each gene. The best model for each euBac protein was selected using IQ-TREE ModelFinder (-m TEST). A maximum likelihood phylogenetic tree was then constructed using the concatenated fasta sequences and the gene partition file, with *Rickettsia canadensis* as the outgroup using the -o option in IQ-TREE. Branch supports were evaluated using 1,000 ultrafast bootstrap approximations (-bb 1000). 92% of the branches exhibited bootstrap >=80% with branches close to root showing >=90% confidence (Fig. S1 D). The overlap between 704 eukaryotes present in the 18S rRNA trees and the *euBac-concat* tree resulted in 423 common organisms. The extra organisms were dropped using drop.tip() function in R resulting in the final *euBac-concat* tree with 423 eukaryotes (51 protists, 152 fungi, 3 choanoflagellates, 43 plants and 174 animals).

Polytomies were removed for all the trees using an inhouse R script. Further, a small value of branch length (10^-6^) was given to branches with branch length <=0. Amongst 704 organisms 96 were protist, 326 were fungi and 61 and 217 belonged to plantae and animalia respectively. This list also includes 4 choanoflagellates All the tree files are provided in Supplementary dataset 4.

In all trees except for the *18S-archaea* tree, the LECA (Last Eukaryotic Common Ancestor) was defined as a single node that overlaps with the root node of the tree. In the case of the *18S-archaea* tree, we defined the LECA as the union of three nodes that appear after the root node and before the nodes showing major diversification among protists.

### Orthologue definition, classification and analysis

To categorise DBDs into orthologous groups (OGs), domain sequences were extracted from the OrthoDB protein sequence list using family position information obtained from Interproscan output. An all-vs-all similarity search was performed for each family using NCBI Blast version 2.5.0 with Evalue cutoff of 10^-3^ (41). Orthologous pairs across 1271 eukaryotic organisms and 1 archaeon for each family were identified using a bidirectional best-hit approach. For each protein in-paralog pairs were identified by selecting only those proteins which show a higher percentage identity than the 75th percentile of the pairwise identity to their respective orthologs. These pairs were then clustered, using pairwise identity as the parameter, into their respective OGs using MCL software version 2.6.3 by using a set of inflation parameters (-I option) for different families (42).

To determine the number of clusters, clustering was initially performed with various inflation parameters (using the -I option), ranging from 0.2 to 5.0. The relationship between the number of proteins and the number of clusters for families that displayed clear trends as the inflation parameters changed was then analysed (Fig. S1 E). For families with 600 or fewer proteins, the maximum number of clusters up to 10 was selected, based on cases where two consecutive inflation parameters resulted in the same number of clusters. For families with more than 600 proteins, the number of clusters was calculated using the slope and intercept from the relationship between the number of proteins and the number of clusters (slope = 0.052997, intercept = -34.2911). Inflation parameters that produced cluster numbers consistent with these calculations and showed saturation in the number of clusters (i.e., where at least two consecutive inflation parameters resulted in the same number of clusters) were then chosen.

This resulted in 34,194 OGs for 704 organisms with 133 TF families and 77 nTF families spanning for 442,013 proteins (Supplementary dataset 5). For the Znf-C2H2 family (IPR013087) proteins, which represents a very large number of sequences, many of which are likely products of duplication being found in tandem in the same protein, a single domain per protein was randomly picked.

To calculate the conservation of OGs for both TF and nTF categories, for each OG, the number of organisms it is present in was counted. Clustering of presence/absence of 34,194 OGs was performed using the “ward” method in clustermap() command from the seaborn module in python. The pairwise euclidean distance for each organism pair was calculated from the OG presence/absence information by using the distance_matix() command from the spicy.spatial module in python. The pairwise_distances() command from the sklearn module in python was used to calculate jaccard distance for all organism pairs and also within kingdom organism pairs.

### Discrete trait reconstruction

The ancestral state for the presence/absence of each OG was estimated using the ace() function from phytools in R, which uses the maximum likelihood method (43). This analysis was performed for the 4 above mentioned phylogenetic trees. Transition rate matrix was established by comparing three evolutionary rate models for discrete states, namely Equal Rates (ER), Symmetric (SYM), and All Rates Different (ARD). The best fit of the model was selected by comparing Akaike Information Criterion (w-AIC) weights and selecting for the maximum value. The discrete state reconstruction of each of the OGs was performed using an in-house R script. An OG was considered to be present at a node if the likelihood value for presence is >=0.70 (Supplementary dataset 6A/B).

### First emergence calculations for OGs

The first emergence point of an OG was defined as the node with the smallest distance from the root node containing that OG. This calculation was performed for each root to tip path using an in-house python script (Suppementary_dataset_7). Using the first emergence information the branch length over which an OG is retained following its first gain was computed using a python script.

To evaluate whether TFs have emerged before nTFs, for each root-to-tip path, the list of TFs and nTF OGs first emerged on that path was obtained and the distance of its emergence from the root calculated (Supplementary dataset 9). The Mann-Whitney statistical test from the python spicy module was used to infer whether TFs have emerged further from the root than nTFs. To calculate the fractions of paths having later emergence of TFs than nTFs the number of such paths having Mann-Whitney FDR <=10^-3^ was computed.

### Gain/Loss calculation

For each node, the total number of OGs present was calculated. For an ancestor-descendent node pair the OGs unique in both descendent and the ancestor were counted. OGs unique to the descendent were considered as OGs gained in comparison to the ancestor and those unique to the ancestor were considered as OGs lost in the descendent. The gain and loss fraction with respect to the ancestor was calculated as the ratio of gain or loss to the total OGs present at the ancestor which is a summation to OGs unique to ancestor and OGs common to both ancestry and descendent (Supplementary dataset 8).

### Simulation

To calculate a random distribution of emergence of OGs across the *euBac-18S* tree, the presence/absence of each OG on the phylogenetic tree was simulated using an in-house R script. To simulate the data, the gain/loss rates obtained from reconstruction of discrete traits for each OG was used. For each iteration the “presence” state to the root was randomly allotted for the same number of OGs as in the reconstructed data. The rTraitDisc() function from the ape package in R was used to predict the discrete state for all the nodes. 100 such simulations were performed.

## Results and Discussion

### Distribution of Transcription factor harbouring superfamilies across eukaryotes

We curated protein superfamilies for DNA-binding domains (DBDs) found in TFs (TF-SF for Transcription Factor-containing Superfamilies). Whereas sequences belonging to the same DNA-binding sequence *family* share strong sequence similarity, members of the same *superfamily* may be divergent at the sequence level but show similarities in their structures and are yet believed to share a common ancestor. Many TF-SFs harbour not only DBDs and families for TFs, but also include non-TF (nTF) proteins. The dataset we had assembled is described in Table 1 and in Supplementary dataset 2.

We identified over ∼880,000 proteins (Table 1, row ‘a’) containing at least one TF-SF across 1271 eukaryotic organisms spanning protists, fungi, plantae and animalia kingdoms. The term protist used in this paper does not refer to a monophyletic clade, but is a term of convenience used to refer to unicellular eukaryotes that do not belong to the fungi, plantae and animalia kingdom. We classified each protein as TF or nTF based on the DBD/sequence family it contains, under the assumption that each family represents a coherent function at this level. We grouped proteins with both TF and nTF DBDs/families as TFs. A small fraction of TF-SF proteins could not be categorised.

TF-SF proteins account for <10% of total coding genes; this ratio increases from unicellular protists and fungi to more complex plants and animals (Fig. 1B). Animalia have the highest number of different TF-SFs whereas protists encode the lowest, consistent with previous findings (19). TF-SFs such as ‘Homeodomain-like*’* (HD) and ‘Zinc finger C2H2’ (ZnF), ‘are highly conserved across all kingdoms consistent with earlier surveys (Fig. 1A) (19, 44). In contrast, a few TF-SFs, such as the ‘Copper-fist DNA binding domain’ in fungi, are uniquely present within certain kingdoms.

The number of TFs exceeds that of nTFs by ∼3-fold (Fig. 1C, Table 1, row ‘b’). Note here that the analyses described here, which directly compare TF and nTF numbers or diversity, were restricted to those TF-SFs containing both TFs and nTFs (Table 1, row ‘b’). The ratio of TF to nTF varies across kingdoms. In the Animalia kingdom, as many as 80% of TF-SF proteins are TFs, whereas in protists and fungi, it can drop to less than 50% (Fig. 1D). Similarly, there is variation in this ratio across TF-SFs. >90% of proteins in the ‘Helix-Loop-Helix’ and “bZip” superfamilies are TFs, whereas <40% for Winged-helix (wH) superfamily are (Fig. S2 A; Fig S1 G and H for variation in HD and ZnF across clades).

Finally, there is a positive relation between the number of TF-SF proteins (Fig S1F), and of TFs and nTFs (Fig. 1E, 1F), and the total protein count (Fig S1 F). We visually observed that different clades of eukaryotes appear to show different relationships (all positive) of TF or nTF count with total protein count. This is unlike in bacteria where most, if not all, genomes together show a single quadratic relationship between the number of TFs and total protein or gene count. Prior literature has shown that the counts of different categories of genes show some power law relationship with the total protein or gene count (45, 46). Keeping this in mind, we assumed that each clade shows its own *y* = *ax^b^* relationship between the number of TFs (*y*) and the total protein count (*x*). We estimated a and *b* for various well-represented clades and found differences in the power between them. For example, whereas *Chordates* show a power of ∼1.6, the *b* for the *Arthropoda* is only ∼0.8 (Fig. 2D). *Streptophyta* (including flowering plants) show a near-linear relationship (*b = 1.01*) (Fig. 2C), as do protists (*b =* 0.95) (Fig. 2A). Among fungi, Ascomycota and *Basidiomycota* show sub-linear powers (*b = 0.64 for*

*Ascomycota and 0.78 for Basidiomycota*); however, the *Mucoromycota* and *Chytridiomycota* are clear outliers showing huge expansions of TFs (Fig. 2B). nTFs show a more or less linear relation for both *Chordata* (*b=0.89*) and *Arthropoda* (*b=1.03*) (Fig. 2H), as do *Streptophyta* (*b*=0.92) (Fig. 2G) and the protists (*b*=0.83, Fig 2E). Fungi exhibit a strong sub-linear relationship for nTFs (*Ascomycota b* = 0.32 and *Basidiomycota b*=0.43). *Mucormycota* and *Chytridiomycota* do not show a huge expansion of nTFs as they do for TFs (Fig. 2F).

**Figure 1:**
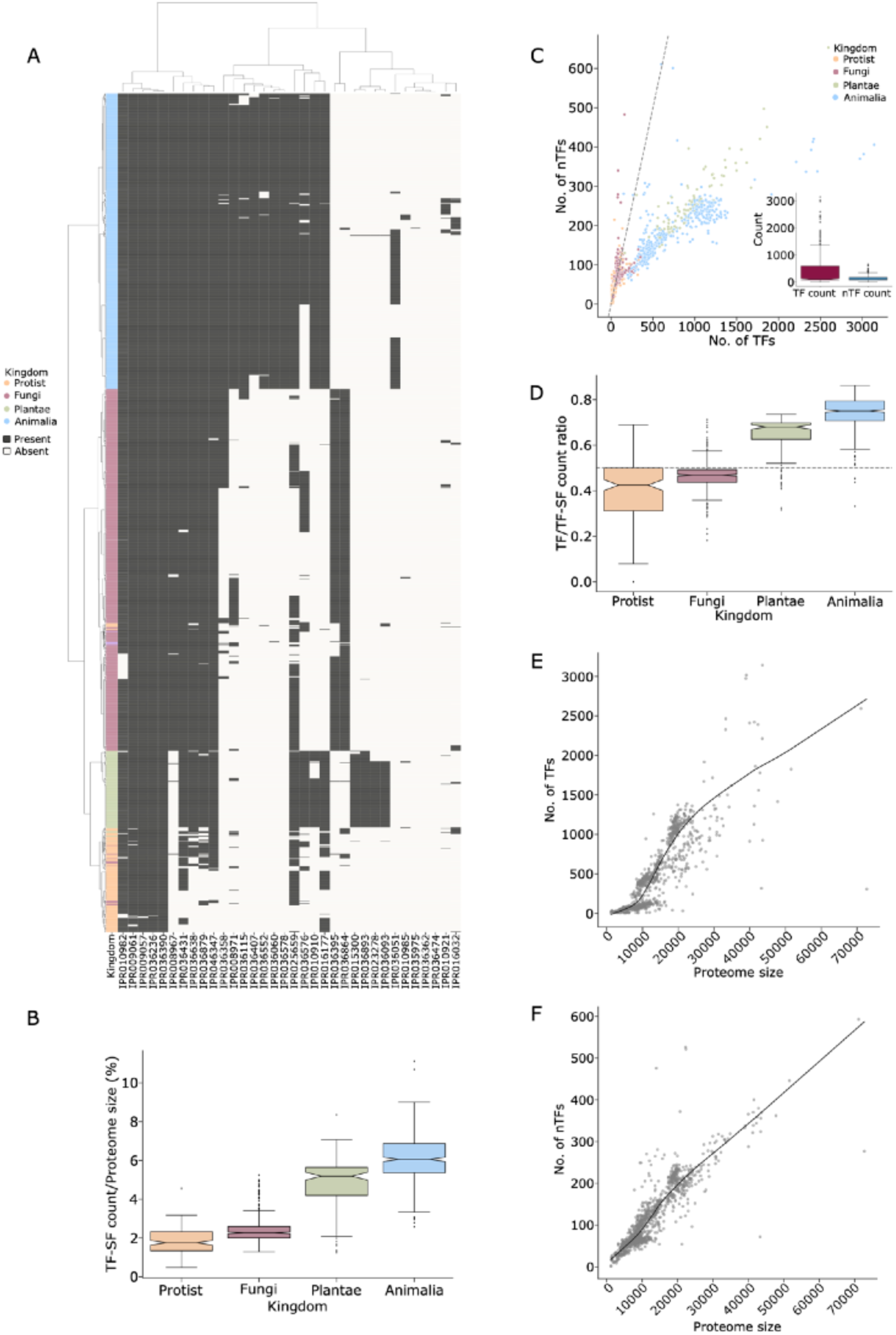
TF and nTF distribution in eukaryotes. (A) The heat map shows the presence / absence of TF-SFs across 1271 eukaryotic organisms clustered using the clustermap() command in python with “ward” as the method. Presence is considered even if a single protein belonging to a TF-SF is present. “Grey” marks the presence and “off-white” marks the absence of a TF-SF. Each column represents a TF-SF (bottom) and each row represents an organism belonging to a kingdom marked on the left. (B) Boxplot displaying the percentage of TF-SF proteins as a fraction of total proteins in organisms across the four kingdoms. (C) Scatterplot of the number of TFs versus the number of nTFs for each organism across the four kingdoms. The dashed grey line marks the 45o line between the x and y axis. The inset shows a boxplot for the number of TFs and nTFs across all organisms. (D) Boxplot displaying the ratio of the number of TFs as a fraction of total TF-SF proteins in organisms across the four kingdoms. (E, F) Scatterplots in figure E and F show the relation between the number of TFs and nTFs respectively, and the proteome size (total number of proteins) for 1271 organisms. The black line represents the regression line between log10 (x) and log10(y) axis as calculated using the HuberRegressor() command in python.

**Figure 2:**
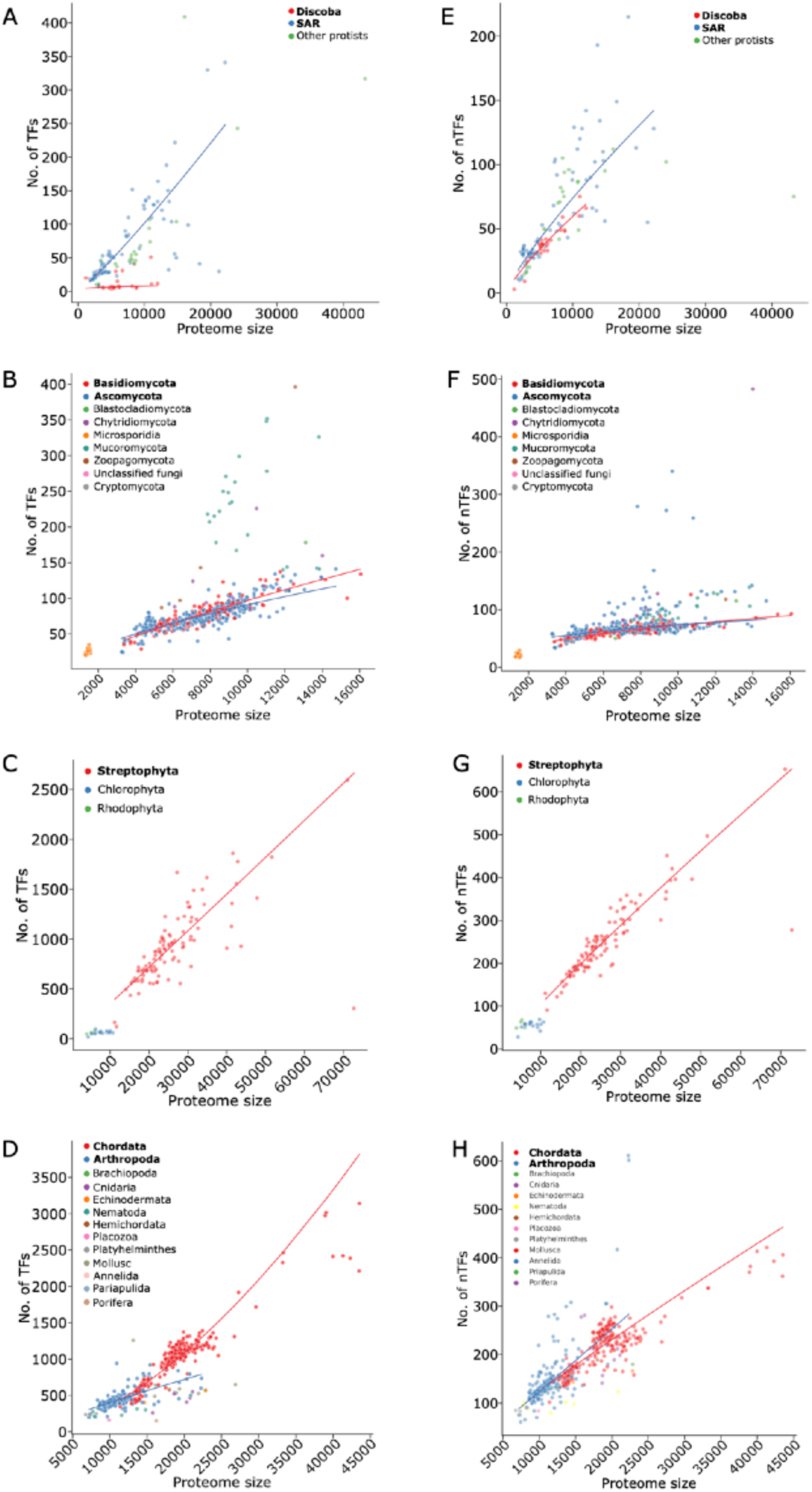
TF and nTF scaling with proteome size. Scatterplot shows the relation between the number of TFs or nTFs and the proteome size (total number of proteins) for different kingdoms. Fig. A,B,C,D represent the TF and proteome size relation for protist, fungi, plantae and animalia respectively. Similarly Fig. E,F,G,H represent the nTF and proteome size relation for protist, fungi, plantae and animalia respectively. For each scatterplot the regression line is plotted for the major phylum within the kingdoms : SAR and Discoba for protists, Ascomycota and Basidiomycota for Fungi, Streptophyta within plantae and Chordata and Arthropoda for animalia. The regression line between log10 (x) and log10(y) axis for each phylum was calculated using the HuberRegressor() command in python.

In summary, whereas overall TF counts far exceed nTF counts, the distribution of TFs and nTFs varies across clades. This is especially so in animalia and plantae. While there is a positive relationship between the number of TFs and nTFs with total protein count, the precise relationship, as determined by the power-law parameter, differs across clades unlike in bacteria.

### State of the last common eukaryotic ancestor(s)

Towards understanding the evolutionary history of TFs and their nTF relatives, we assembled multiple phylogenetic trees for eukaryotes (Supplementary dataset 4). We first built an alignment of 18S rRNA sequences from 704 genomes. To build phylogenetic trees we took the following approach. Two recent papers have proposed two different topologies for the eukaryotic tree, each driven by a different root. Based on alignments of eukaryotic genes of bacterial origin, Derelle et al. (PNAS 2014) proposed a root between the branches leading to *Opimoda* (*Archaeaplastida*, SAR) and *Diaphoda* (*Opisthokonta, Amoebozoa*) (35). In contrast, using genes of archaeal origin, Jewari et al. (Science 2023) more recently proposed an “*Excavata*” root for the tree (36). We used these tree topologies as constraints on our 18S rRNA alignment to build two trees, henceforth referred to as “*euBac-18S*” and “*euArch-18S*” trees (Fig. 3A,B). These two trees differ considerably in the branch length separating the root from the *Opisthokonta* common ancestor and *Streptophyta* common ancestor, with the *euBac-18S* tree placing the root much closer to these clades (*Opisthokonta* branch length: 0.011, *Streptophyta* branch length: 0.02315) than the *euArch-18S* (*Opisthokonta* branch length: 0.053, *Streptophyta* branch length: 0.0596). Note that the *Opisthokonta* include the *Mammalia* and *Streptophyta* includes flowering plants, both of which code for the largest numbers of TF-SF proteins on average in our dataset. We also built an unconstrained 18S rRNA tree, rooted using a 16S rRNA sequence from an archaeon *Metallosphaera sedula*, referred to as the “*18S-archaea*” tree (Fig. 3C). We also used an unconstrained tree comprising 423 eukaryotic genomes, which was constructed by concatenating the sequences of 25 euBac proteins and rooted using a presumed alpha-proteobacterial ancestor of the mitochondria; this we call the “*euBac-concat*” tree (Fig. 3D).

**Figure 3:**
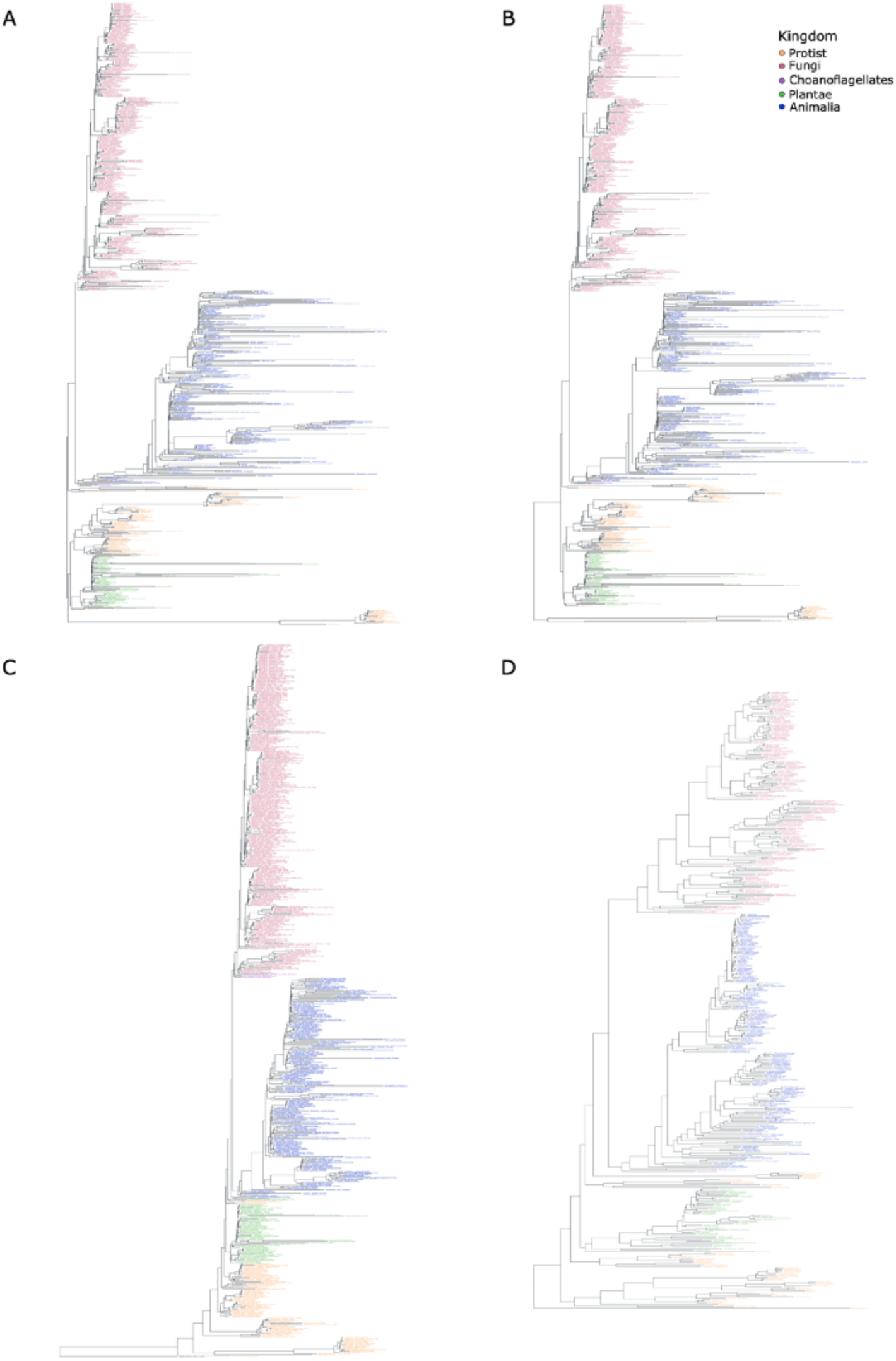
Tree topologies. The figure shows the different tree topologies used in the study. (A) euBac-18S tree with 704 eukaryotes made using the tree provided by Derelle et al. (PNAS 2014) as constraints on the 18S rRNA alignment. The tree is rooted at the clade similar to the constraint tree. (B) euArch-18S tree with 703 eukaryotes made using the tree provided by Jewari et al. (Science 2023) as constraints on the 18S rRNA alignment. The tree is rooted at the clade similar to the constraint tree. (C) 18S-archaea tree with 704 eukaryotes and 1 archaeon (Metallosphaera sedula) as the outgroup. The tree was made using 18S rRNA sequences obtained from the SILVA database. SILVA ACT online web server was used to make the tree under GTR as the evolutionary model. (D) euBac-concat tree with 423 organisms was made using orthologs of 25 eukaryotic genes of bacterial ancestry (euBac genes He et. al 2014). The tree was rooted using Rickettsia canadensis as the outgroup.

**Table 2:** TF and nTF repertoire in LECAs.

| Tree | No. of TF OGs | No. of nTF OGs | No. of UN OGs | No. of TF family | No. of nTF family | No. of UN family | No. of TF-SFs |
| --- | --- | --- | --- | --- | --- | --- | --- |
| *euBac-18s* | 599 | 342 | 0 | 19 | 41 | 0 | 12 |
| *euArch-18s* | 100 | 108 | 0 | 18 | 32 | 0 | 12 |
| *18S-archaea* | 42 | 33 | 0 | 7 | 20 | 0 | 8 |
| *euBac-concat* | 509 | 260 | 1 | 22 | 44 | 1 | 12 |

**Table 3:** Overlap between TF repertoires of LECAs.

| Tree1 | Tree2 | category | OG | Family | TF-SF |
| --- | --- | --- | --- | --- | --- |
| *euBac-18s* | *euArch-18s* | tf | 94 | 18 | 11 |
|  |  | ntf | 92 | 32 | 5 |
| *euBac-18s* | *18S-archaea* | tf | 17 | 7 | 6 |
|  |  | ntf | 12 | 20 | 4 |
| *euBac-18s* | *euBac-concat* | tf | 187 | 19 | 11 |
|  |  | ntf | 91 | 39 | 6 |
| *euArch-18s* | *18S-archaea* | tf | 13 | 7 | 6 |
|  |  | ntf | 9 | 18 | 4 |
| *euArch-18s* | *euBac-concat* | tf | 42 | 18 | 11 |
|  |  | ntf | 41 | 32 | 5 |
| *18S-archaea* | *euBac-concat* | tf | 36 | 7 | 6 |
|  |  | ntf | 22 | 20 | 4 |

We then used a bidirectional best hit approach followed by network clustering to group ∼800,000 TF-SF family sequences from 704 eukaryotic genomes into 34,191 orthologous groups (OG). We performed this at the family level rather than the whole protein sequence level to cover for the possibility of distinct evolutionary trajectories of individual domains/families in a multidomain protein. We forced each OG to be populated by the members of the same sequence family or domain and therefore the same function - TF or nTF - resulting in 28,493 TF and 5,693 nTF OGs. The presence / absence matrix of OGs loosely clusters organisms by their kingdom (Fig. S2C). There was little difference between TF and nTF OGs in the extent to which they are conserved across genomes in this study (Fig. S2B).

Considering each OG as a unit of evolution, we reconstructed its presence or absence on the tree using maximum-likelihood (Supplementary dataset 6B). We first examined the repertoire of TF-SF OGs in the last eukaryotic common ancestor (LECA) for each phylogenetic tree. For each LECA (one for each tree), we catalogued the presence of both transcription factor (TF) and non-transcription factor (nTF) OGs. Despite differences in OG counts in the LECA across the trees (Table 2, Supplementary Table 1), some consistent patterns emerged. Some families of TFs, such as TFIIIB, bZIP, Myb_domain, AP2/ERF domain, ZnF-C2H2-LYAR-type, Tubby, Cro/C1-type helix-turn-helix, were present in the LECA across trees. The nTF families in LECA include multiple chromatin remodeler families, enzyme families and ribosomal protein families. Finally, the overlap, in the OGs and families present in the LECA, between the two constrained trees, which we consider as the best available hypotheses for the correct tree rooting, is considerably high (Table 3).

We compared our TF predictions for the LECA with two previously published studies. Iyer et al. (2008) (44) examined 55 superfamilies of DNA binding domains and concluded that 15 TF families were either present in LECA or emerged very early in eukaryotic evolution. Our analysis shows that 7 of these TF families overlap with the *euBac-18S* and *euArch-18S* trees. These overlapping families include bZIP, Myb, AP2/ ERF, Heat shock protein, Forkhead, Znf-C2H2, and E2F/DP. While Iyer et al. also reported a bHLH family at LECA, our inferences do not include the same bHLH family but do identify Myc-bHLH, a member of the bHLH superfamily. We also compared our results with those of Alex De Mendoza et al. (2013) (19), who used a Dollo parsimony-based phylogenetic approach to predict LECA’s TF families. Our inferences match 10 out of the 27 TF families they had identified.

Among extant OGs, the ratio between TFs and nTFs is 3.8. However, that in the LECA, considering OGs conserved in the LECA in both the constrained trees, is ∼1. The highest TF:nTF ratio we saw among the 18S trees is 1.8 in the *euBac-18S* tree, in which the root is placed closer to the “higher” eukaryotes than in the other trees. The TF:nTF ratio in the LECA of the *euBac-concat* tree, which shows a strong overlap with the LECA of the *euBac-18S tree,* is nearly 2, but significantly less than that expected from the extant genomes in our dataset (*P* < 10^-10^ for comparisons of each LECA with extant, Fisher’s Exact test).

Thus, though the LECA comprised tens to hundreds of TF and nTF OGs, pointing to the presence of a complex regulatory and chromosome organisation system, it probably coded for an under-representation of TF OGs, compared to the average across all extant genomes in our dataset.

### Evolutionary dynamics of TF-SFs

To understand the dynamics of evolution of TF-SF DBDs across the eukaryotic phylogenies we used our maximum likelihood estimation of presence/absence state of OGs in several ways. For each branch, we identified the set of OGs gained or lost in the descendent node in comparison to the immediate ancestor. We found that between 66-74% of such gains are retained all the way to the tip along at least one downstream path, across the 4 phylogenies. These OGs have spent significantly more time along the corresponding path when compared to those showing a loss following the gain (P<10^-15^ for all trees, Mann-Whitney). Overall, the average TF-SF OG is retained over a maximum of ∼20% of the mean branch length on the tree (Fig. 4A,B, Fig. S3 A,B).

Between TFs and nTFs, nTFs seem to retain a slightly larger fraction of such gains up to the tips than TFs (81% for nTFs and 71% for TFs); this difference is statistically significant (P<10^-15^ and mean odds ratio 0.572, Fisher’s Exact test) (Fig. 4A,B, Fig. S3 A,B). TFs and nTFs also show a statistically significant difference between their overall retention distances across all trees, with nTFs being retained over longer branch lengths than TFs (P<10^-15^ for all trees, Mann-Whitney); the absolute difference however is quite small (Fig. 4A,B, Fig. S3 A,B). We also observed that TFs retained at the tips were gained more recently than equivalent nTFs (P<10^-15^ for all trees, Mann-Whitney). Thus, TFs appear to be retained over shorter evolutionary distances than nTFs across all trees.

**Figure 4:**
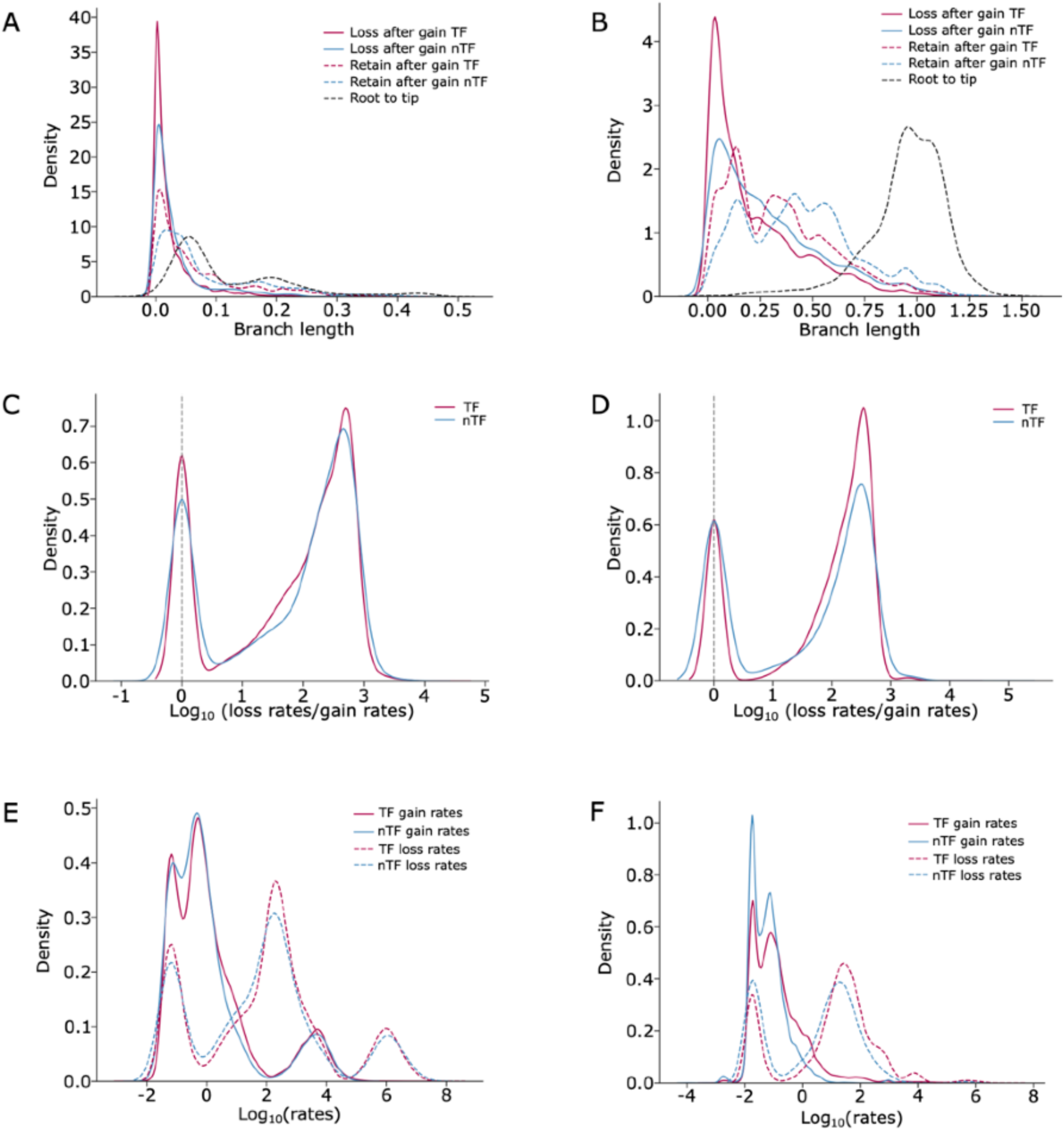
Gain and loss trends of TFs and nTFs. (A) The density plot depicts the branch length distribution for the length travelled by an OG after being gained for the first time for (A) euBac-18S and (B) euBac-concat trees. The dark pink colour is for TFs and blue is for nTFs. The branch length for loss of OG after its first emergence is denoted by solid lines for both TFs and nTFs. Similarly, branch length for OG being retained after its first emergence is denoted by dashed lines. The black dashed line represents the overall distribution of branch length from root to tip. Refer to figure S3 A and B for euArch-18S and 18S-archaea trees. Figure C and D represent the distribution of log10 of loss rates/gain rates for each OG for euBac-18S and euBac-concat trees respectively. The dark pink colour is for TFs and blue is for nTFs. The vertical grey dashed line depicts the value where loss rates and gain rates are equal. Refer to figure S3 C and D for euArch-18S and 18S-archaea tree. Figure E and F represent the distribution of log10 transition rates for gain (from absence to presence) and loss (from presence to absence) of OGs for euBac-18S and euBac-concat trees respectively. The solid lines refer to gain rates for TF (dark pink) and nTFs (blue). The dashed lines refer to loss rates for TF (dark pink) and nTFs (blue).

Consistent with the short retention time of OGs, we found that the rate of loss, as measured by maximum likelihood parameters estimated for each OG during ancestral reconstruction, is greater than that of gain for most OGs for both TFs and nTFs (Fig. 4C, D, Fig. S3 C,D) (Supplementary dataset 6A) . This is true for all the phylogenies considered. TFs exhibit slightly but significantly higher rates for both gain and loss than nTFs (*gain rate* P< 10^-6^ and *loss rate* P < 10^-8^ for all trees, Mann-Whitney) (Fig. 4 E,F, Fig. S3 E,F). Whether TFs and nTFs show differences in the *loss rate*: *gain rate* ratio, however, seems highly dependent on the tree used.

For each of the nodes across the four phylogenies we then calculated the number of OGs gained or lost (number of OGs gained or lost as a fraction of that in the ancestor) in each descendant (Supplementary dataset 8) (Fig. S4). Independent of the phylogenies, the overall fraction of OGs lost exceeded the gain fraction for all trees and was statistically significant different for *euBac-18* (P=0.009, Mann-Whitney)*, 18S-archaea* (P=0.002, Mann-Whitney) and *euBac-concat* (P< 10^-15^, Mann-Whitney) trees. Between TFs and nTFs, nTFs consistently exhibited more loss than gain across trees (P < 10^-6^ for all trees, Mann-Whitney). For TFs, these numbers were inconsistent across trees (Fig. S4). At the kingdom level as well, for both TFs and nTFs, the gain:loss ratio varies considerably across trees (Fig. S4).

In summary, nTFs appear to be retained over slightly longer evolutionary distances than TFs. The evolution of TF-SFs is mostly driven by higher loss rates than gain. In general, the overall flux of TFs, measured by gain or loss rates, is slightly higher than that of nTFs. The loss/gain ratio, in particular the question of whether this value significantly differs between TFs and nTFs and across clades is highly sensitive to the topologies of the tree in the multicellular clades.

### Expansion of TFs and nTFs

We have shown that TF-SF OGs are dynamic over the course of evolution and are retained, on average, ∼20% (across the four trees) of the mean evolutionary distance from root to tip. We now asked where different TF and nTF OGs were first gained along an evolutionary path, and whether TF and nTF OGs differ from each other in this respect. For this, for each path on the phylogenetic tree (from root to tip), we calculated the first emergence of each OG on that path. For every OG we then calculated the number of unique nodes, computed across paths, at which it was first gained. We observed that across the four phylogenies 57-73 % of OGs show a single node of emergence across all paths (Fig. 5A inset, Fig. S5 A,B,C). To test how these numbers would differ from null expectations, we simulated the evolutionary trajectory of each OG on the *euBac-18S* tree, given a random initial state (see Methods) and maximum likelihood gain and loss rates estimated from the presence/absence data for extant species. In these simulations, on average, only 35% of OGs were inferred to have been gained once (Fig. 5A). This difference between simulated and observed values is statistically significant (P<10^-15^). Thus, a majority of TF and nTF OGs were gained just once along the phylogeny, and this is not merely a product of its gain and loss rates along the phylogeny.

**Figure 5.**
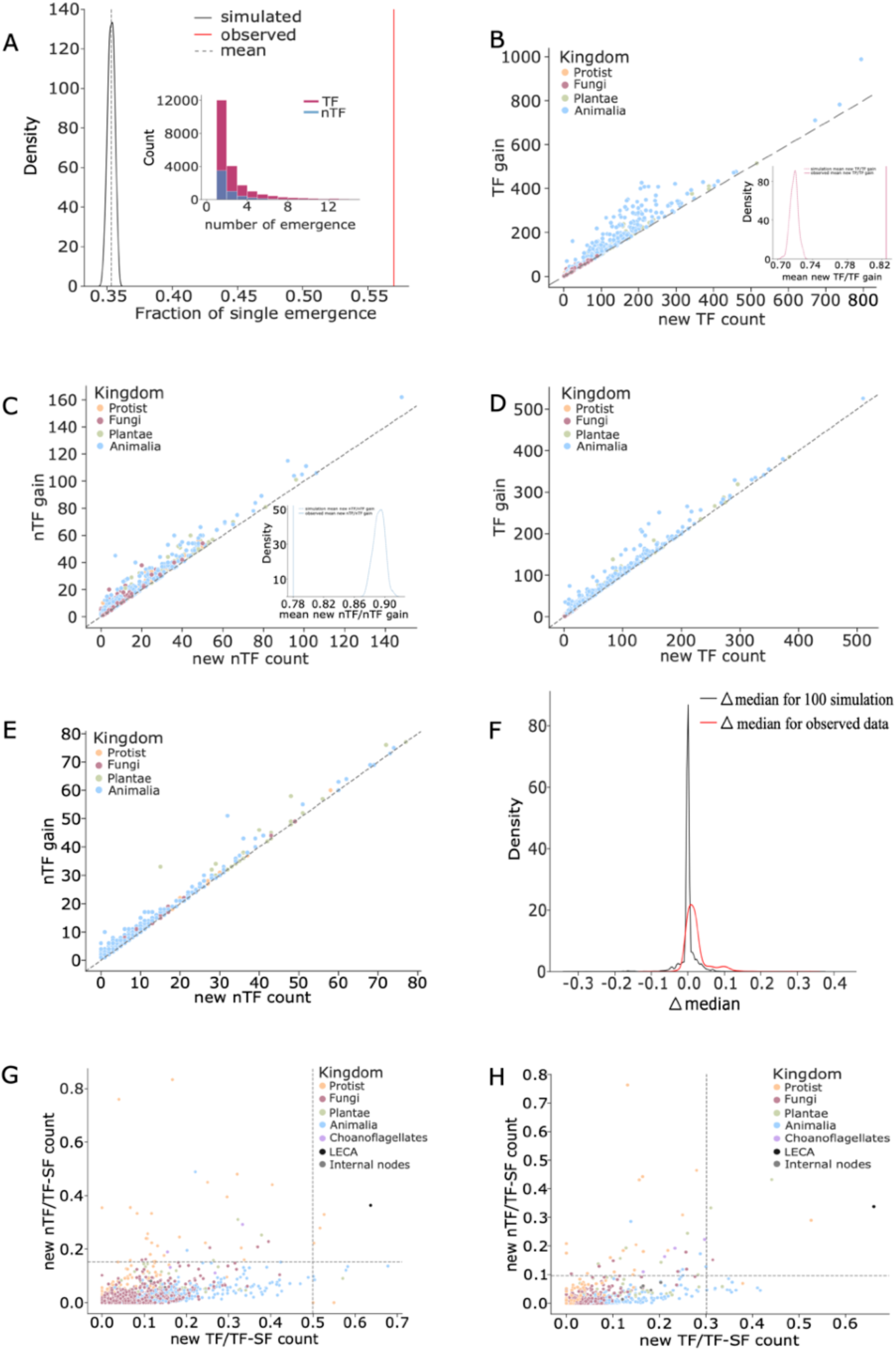
Emergence statistics of TFs and nTFs. Figure A represents the distribution of the fraction of OGs exhibiting a single instance of emergence. The distribution in black represents this fraction for the simulated data with a grey dashed line depicting the mean of the distribution. The line in red represents the observed ratio for the euBac-18S tree. The inset shows the histogram for the number of independent emergence for TF (dark pink) and nTF (blue) OGs for the euBac-18S tree. Here we show a maximum of 14 instances of emergence but it is to be noted that the value goes upto more than 100 emergence for certain OGs. Refer to figure S4 A, B and C for the distribution of the number of emergence for euArch-18S, 18S-archaea and euBac-concat trees. Scatterplot for gain of new OG versus emergence for new OG at a node was plotted for both TFs and nTFs across the four kingdoms. Figure B and C depict this relation for the euBac-18S tree for TF and nTF respectively. Figure D and E depict this relation for the euBac-concat tree for TF and nTF respectively. The dashed grey line marks the 45o line between the x and y axis. The inset within Figure B shows the distribution of new TF to gain TF ratio obtained from the simulations (dark pink dashed line) and the observed value (dark pink solid) for the same for euBac-18S tree. The inset within Figure C shows the distribution of new nTF to gain nTF ratio obtained from the simulations (blue dashed line) and the observed value (blue solid line) for the same for euBac-18S tree. Refer to figure S4 D and E for euArch-18S and S3 F and G for 18S-archaea tree. Figure F shows the distribution of the difference between the median TF emergence and nTF emergence (delta median) from the root for each path (root to tip) for the euBac-18S tree. The distribution in the black represents the delta median value for all the paths across 100 simulations and the one in red shows the observed delta median for the euBac-18S tree across all paths. The scatterplots in figure G and H shows the relation between the ratio of new nTF to total TF-SF proteins to the ratio of new TF to total TF-SF proteins for both internal node and tips for the euBac-18S and the euBac-concat tree respectively. The vertical grey line marks the Q3 + 3 * IQR value (outliers for the distribution) for the x axis (TFs) and the horizontal grey line marks the same for y axis (nTFs). Refer to figure S4 H and I for euArch-18S and 18S-archaea trees.

At least 87% of the gains across all nodes, when compared to their immediate ancestor, are a result of new innovations, these being the first appearance of the OG along the evolutionary path under consideration (median for TFs across all trees: 88-94% and for nTFs: 87-98%) (Fig. 5B, C, D, E Fig. S5 D,E,F,G). This number appears to be larger than expected from simulations on the trees for TFs (mean across simulations ∼72%, Fig. 5B inset) but not for nTFs for which the mean across simulations (∼89%, Fig. 5C inset) is in fact slightly larger than what we observe.

We evaluated if the distance from root at which a TF first emerged, on average, differed from that at which an nTF emerged (Supplementary dataset 9). We observed that 43-59% of all evolutionary lineages or paths from root to tip across the four trees show an earlier emergence of nTFs than TFs (at FDR <10^-3^). We did not observe any path exhibiting earlier emergence of TFs than nTFs. We also calculated the same for our simulations. We found that none among 100 simulations showed a higher fraction of paths with TFs emerging later than nTFs than our observed data (Fig. 5F). Thus, consistent with the statistical over-representation of nTFs in LECA, TFs on average have emerged later than nTFs in a substantial proportion of evolutionary trajectories. This difference between TFs and nTFs in their node of emergence appears to be driven by two large superfamilies, the ZnF and the wH. The third large superfamily, the HD, however, shows no such relationship.

**Figure 6:**
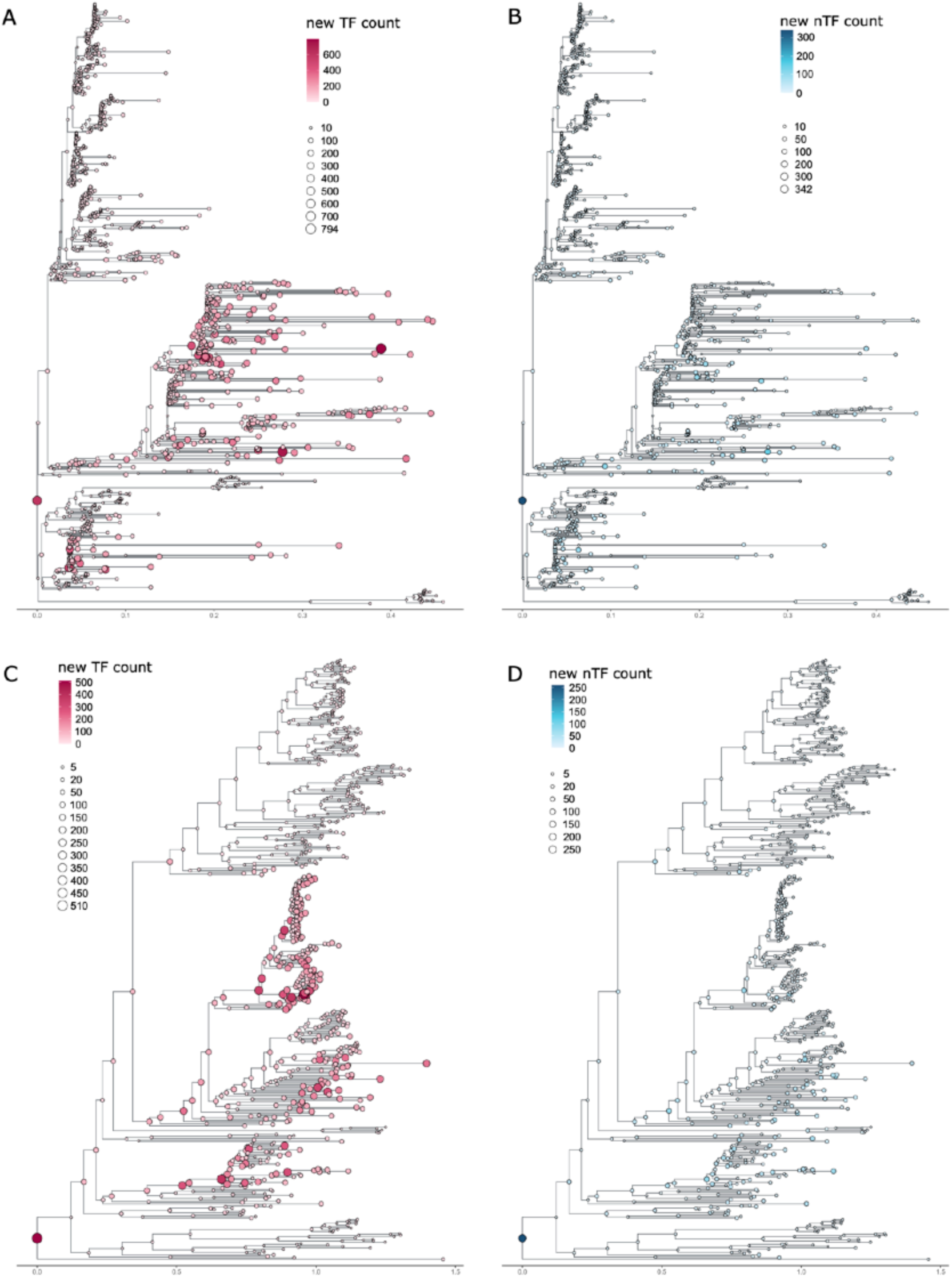
Emergence of TFs and TFs on trees. The figure shows the number of innovations (first emergence) of OGs occurring at each node on the phylogenetic tree. Figure A and B depict the number of innovations for the euBac-18S tree for TF and nTF respectively. Figure C and D depict the number of innovations for the euBac-concat tree for TF and nTF respectively. The dark pink colour represents the TF and blue represents nTF. The darker shades for both TF and nTF represent high values of innovations at a node and lighter shades represent low values. The same goes for the size for the bubbles with bigger size bubbles exhibiting high values and smaller bubbles showing low values. The topologies for the trees are the same as mentioned in the figure. 3 A and D respectively for the euBac-18S and the euBac-concat tree. Refer to figure 6 A and B for euArch-18S and S3 C and D for 18S-archaea tree.

Finally, we visually observed that a few nodes on the phylogeny show large bursts of new TF OG appearances (Fig. 6, Fig. S7) (Supplementary dataset 10). We systematically defined nodes showing bursts of OG innovations as outliers of the distribution of the number of new TF appearances normalised by the total TF-SF OG count for the node (>= Q3 + 3 * IQR) (Fig. 5G, H, Fig. S5 H, I). This resulted in 12-23 of outlier nodes for TF emergence and 41-64 nodes for nTFs across the trees. A large number (9-16 for TFs and 23-31 for nTFs) of these nodes are tips rather than internal nodes. Most of the TF outlier nodes belong to multicellular clades like *Chordata, Arthropoda* and *Streptophyta*. This was in contrast to the nTF outlier nodes, which are populated by mostly various protist and fungal clades. This difference between TFs and nTFs in outlier nodes also presents an explanation for why TFs appear to emerge further from the root than nTFs.

We assessed the contribution of these nodes showing bursts of TF emergences to OGs present in downstream extant organisms. 18-34% of TFs present in the extant organisms arose in these small numbers of outlier nodes showing large numbers of new TF appearances. We had also noted earlier that certain clades such as the animalia show high TF:TF-SF or TF:nTF ratios. We asked to what extent these outliers, showing bursts of TF innovations, are responsible for producing these high TF:TF-SF ratios (TF:TF-SF ratio>=0.65, Fig. S6). For this we looked at the extant nodes with high TF:TF-SF ratio (276-277 extant nodes in the 18S rRNA trees and 207 nodes in the euBac-concat tree) and asked what fraction of these nodes are downstream of TF outliers. We found that for *euArch-18S* and *18S-archaea* trees more that 93% of such extant nodes lie downstream of TF outliers (*euArch-18S*-256/276 and *18S-archaea*-269/277 nodes, (P < 10^-10^, Fisher exact test). For the other two trees this fraction was comparatively low but significant (*euBac-18S*-136/276 and *euBac-concat*-47/207 nodes, P <10^-23^, Fisher exact test). This low fraction is mostly influenced by the TF outlier cutoff, which might exclude some borderline internal nodes. For instance, for the *euBac-concat* tree if the outlier cut-off is decreased to >= Q3 + 2 * IQR, the fraction increases to 99% (205/207 nodes, P < 10^-14^, Fisher exact test). This indicates that high TF counts in these organisms are significantly influenced by burst new TF OG appearance and are not a result of gradual increase in the TF proteins.

In summary, a large majority of TF-SF OGs exhibit a single node of emergence on the phylogenetic tree. Further, along a substantial fraction of evolutionary paths, nTFs from the wH and the ZnF superfamily have emerged closer to the root TFs, with the reverse being rare. TF expansions appear to be driven by a few nodes showing the first emergence of a large number of TF OGs.

## Discussion

In this study, we have described the evolutionary trajectories of DNA binding superfamilies present in eukaryotic TFs (TF-SFs). These TF-SFs include families associated not only with TF function but also with more diverse non-TF roles such as chromatin-remodelling, enzymatic activity, RNA binding roles, DNA repair etc. Different clades of eukaryotes show different scaling of TFs (and nTFs) with genome size, with chordates showing a high power of >1.7 while the number of TFs in fungi scale sub-linearly with genome size. This super-linear relationship in chordates is not reflected for nTFs. This might be consistent with the idea that increase in the so-called organism complexity, defined by structural as well as developmental complexity, might be associated with an excess of regulatory functions and less so with overall numbers of protein-coding genes in higher eukaryotes. Our analysis does not include other types of regulatory mechanisms, the most prominent among these being post-transcriptional control by small non-coding RNA, whose extensive deployment also appears to be linked to this complexity (47). This does not easily translate to bacteria however, where a single near-quadratic relationship fits the scaling of TF counts with genome size or the number of proteins encoded by the genome; presumably, complexity in bacterial lifestyles scales linearly with number of proteins or genes in a genome, unlike in eukaryotes.

Most TF-SFs appear to have been represented in the LECA, irrespective of the tree topology or its rooting. Thus, the LECA already coded for a complex regulatory system, consistent with previous reports. Nevertheless, relative to extant clades, TFs appear to be under-represented, when compared to nTFs, in this ancient organism. This issues with the caveat that presence of a particular protein at LECA does not necessarily imply that it is performing the same function as in extant beings.

TFs and nTFs belonging to TF-SFs are dynamic in being gained and lost frequently. Both TFs and nTFs last only about 20% of the average root-tip distance on the tree. For both TFs and nTFs, losses seem more common than gains. At first glance, this seems to contradict a previous report by Schmitz et. al. (2016) in which TF evolution was shown to be driven by gain (21). However, whereas Schmitz et al. defined TFs by their entire domain architecture, our study operates at the level of individual sequence domains and families. Further, our study differs from Schmitz et al. in how loss is defined. In our study, loss and gain are symmetric: every gain or loss of an OG is counted. In contrast, Schmitz et al. defined loss by the complete disappearance of an entire protein group (which they call DAC), while addition of even a single member of a DAC counted towards a gain. This predominance of loss is similar to results obtained across gene functions in bacteria(48). Though eukaryotic genomes appear to grow in size extensively, genome growth is often linked to selfish genetic elements, which contribute little to our analysis. In fact even for mammals, analysis of pseudogenes suggest that DNA loss may exceed that of gain, again similar to results for pseudogenes obtained in bacteria (49, 50). TFs appear to be subject to greater gain and loss fluxes than nTFs, being gained and lost more frequently than nTFs. Previous studies in different organisms had suggested that regulatory proteins, often defined by TFs, are less well conserved than non-regulatory proteins (13, 51). Though we see little difference in conservation between TFs and nTFs, the higher fluxes for TFs along the phylogeny supports this view.

Gains of TFs and nTFs appear to be driven by a few nodes showing large innovations of new sequence clusters. TFs expanded predominantly in the multicellular clades of *Chordata* and *Streptophyta*, with a notable expansion being the *Mucoromycota* clade within fungi, whereas nTFs showed expansions within different clades of protists and fungi. It has been suggested previously that TF expansion in the multicellular clades, especially within metazoan, is driven by multiple whole genome and segmental duplications events (21). In particular, the huge expansion of TFs in the common ancestor of chordates may be driven, in part, by whole genome duplication. To what extent this explanation holds depends on whether sequence divergence post-duplication is large enough for duplicates to be split across multiple OG clusters in our analysis. We do not address this, or the mechanism by which expansions have occurred in TF and nTF repertoires, in this study. Nodes exhibiting bursts of TF innovations seem to drive the TF content in their descendants with upto ∼30% of the TFs within these organisms emerging at these small number of outlier nodes. Expansion of TFs and nTFs seem somewhat uncoupled for a large proportion of extant organisms with high TF:nTF ratios seem to be descendants of nodes with large numbers of new TF appearances.

Given the early emergence and innovations within nTFs as compared to TFs and the high sequence specificity of TFs, an obvious question that emerges is that specificity might have evolved from non-specific DNA binding families. In other words, might TFs have evolved from nTF-like ancestors, as proposed in an opinion article by Busby and Visweshwaraiah(4)? We don’t test this hypothesis due to limited data at the family level. But under the assumption that members within a TF-SF share a common ancestor, the evidence of early emergence of nTFs than TFs from two large TF-SFs (wH and ZnF) makes these superfamilies well suited candidates to systematically answer this question. That said, prokaryotes make a better group of organisms to address the question of the very emergence of TFs, as LECA is already a late-emerging complex organism.

While our analysis is robust across multiple phylogenies, it is constrained by the assumption that each family represents a coherent group of TFs and nTFs. This assumption may not be entirely valid, for some mutations can potentially alter the binding specificity thereby changing a TF to nTF and vice versa without changing its family. An example for this is the MYB/SANT family within the Homeodomain superfamily, where a difference of few residues makes MYB a TF and SANT a chromatin remodeler (nTF). This assumption however is probably the best that can be made today for a study on this present scale. Our analysis of the evolution of TFs in eukaryotes, as defined by the presence or absence of TF DNA binding families, covers one part of the regulatory network, and how changes in DNA binding specificity of conserved TFs change the structure and function of the regulatory network and determine new phenotypes is an exciting and active avenue for research (52, 53).

## Acknowledgements

We thank Sunil Laxman and Dasarathi Palakodeti for discussions on the project. We specially thank Sunil Laxman for his valuable inputs on the initial draft of the manuscript. We would also like to show our gratitude to Mohak Sharda, Vairavan Lakshmanan and Nitish Malhotra for their valuable inputs.

## Data availability

All supplementary datasets are available on Figshare (https://figshare.com/s/4999f078009c84ea67b1)

