## Supplementary Figures for "Evolution of transcription factor-containing superfamilies in Eukaryotes"

### Figure S1

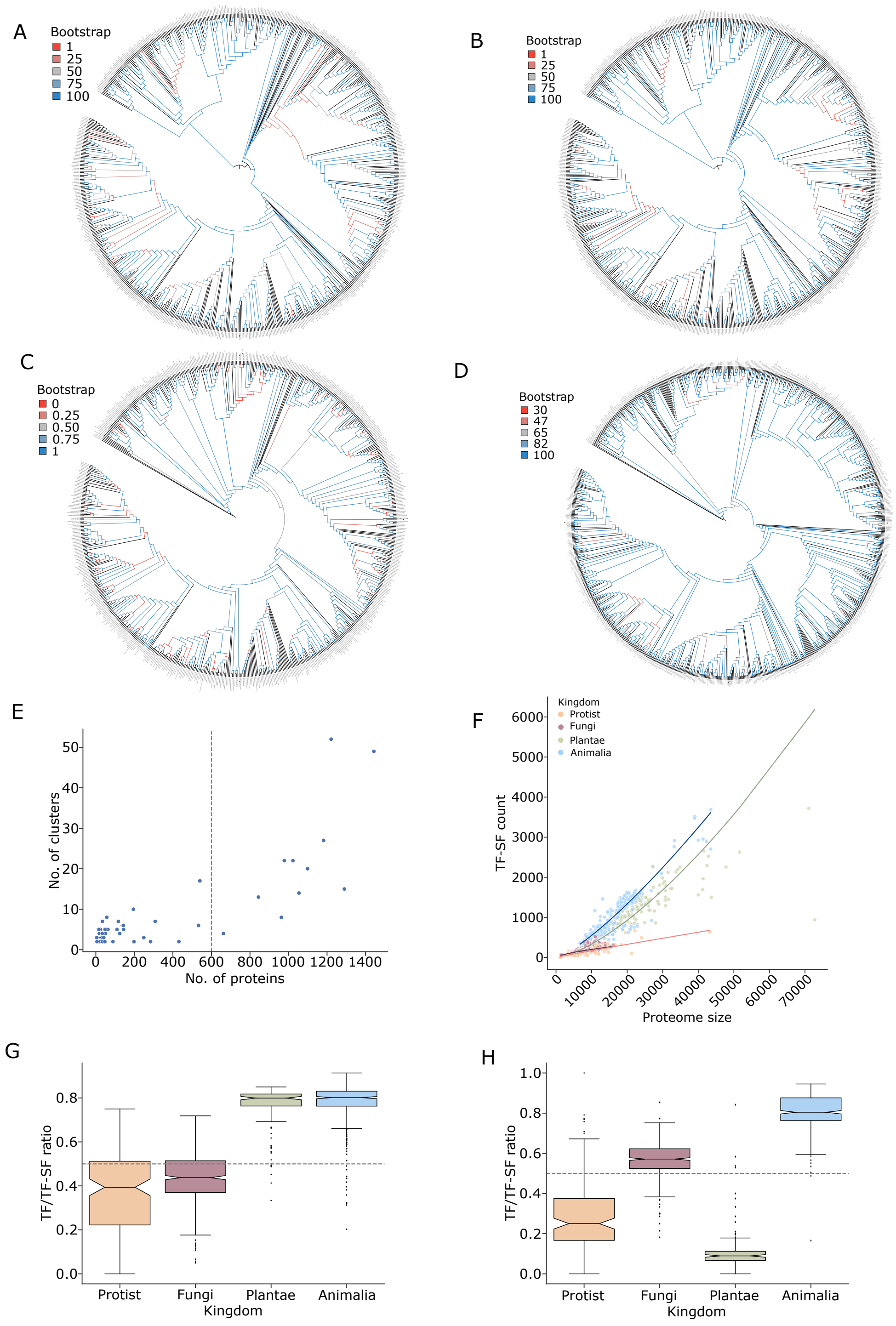


Figures A,B,C and D show the bootstrap values for each branch for the euBac-18S (704 organisms), euArch-18S (703 organisms), 18S-archaea (705 organisms) and euBac-concat (869 organisms, before pruning) tree respectively. It is to be noted that for each tree the high bootstrap values are denoted by blue and the low values by red.

Figure E shows the relation between the optimum number of clusters and the number of proteins. The number of clusters is obtained as a result of varying the inflation parameter, where at least two consecutive inflation parameters resulted in the same number of clusters and the one with the lower value of inflation parameter is considered. The dashed grey line is placed at the number of proteins equal to 600 for a family.

The scatterplot in figure F shows the relation between the total number of TF-SF proteins and the proteome size across the four kingdoms. The solid lines represent the regression line between x and y axis as calculated by the HuberRegressor() command in python.
Figure G and H represent the ratio of TF to total TF-SF proteins for Homeodomain and ZnF-C2H2 superfamilies respectively across the four kingdoms. The horizontal dashed grey line denotes the value where 50% of the proteins within each TF-SFs are TFs.

### Figure S2

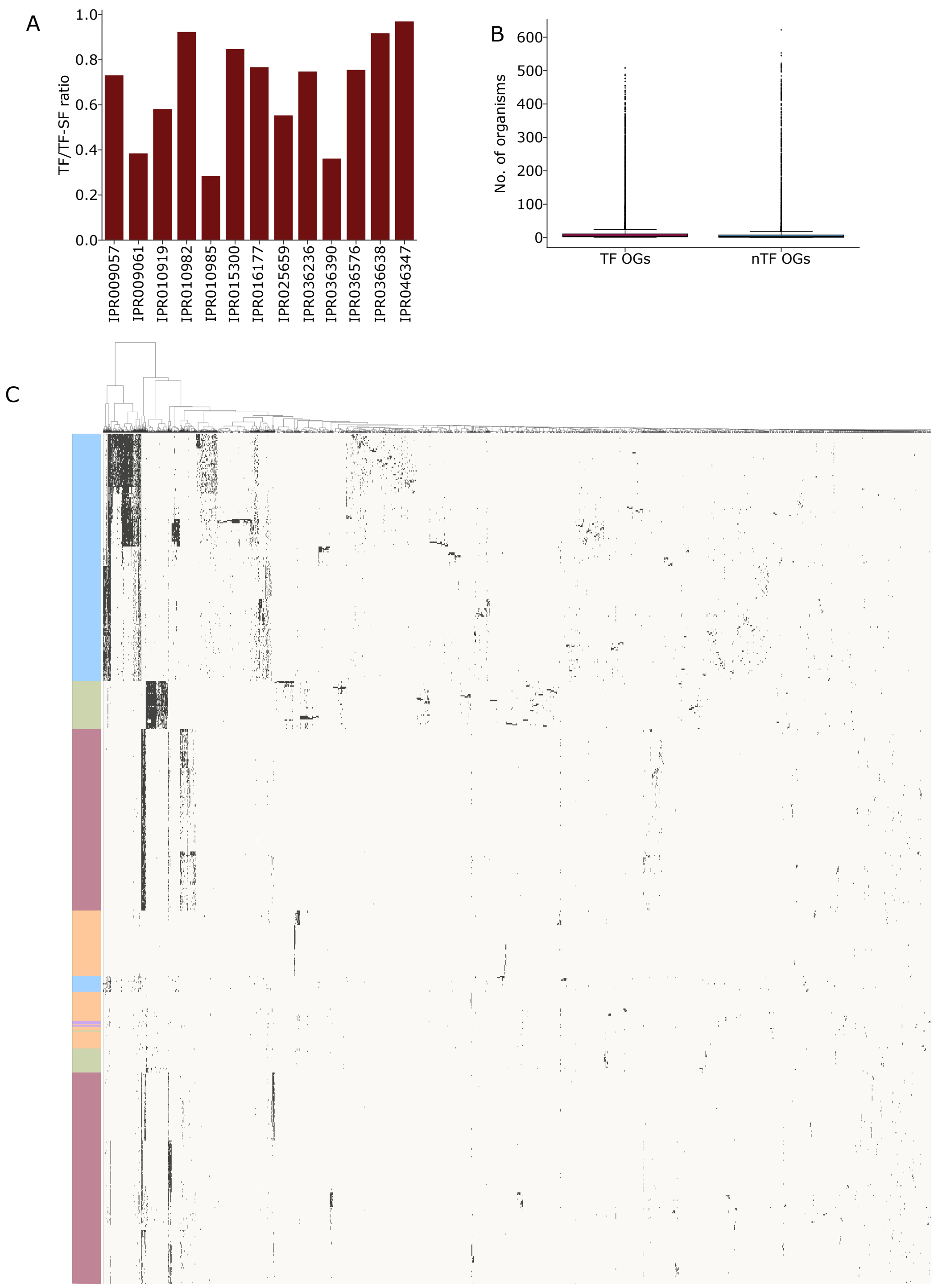


Figure A shows the ratio of total TFs across all organisms to total TF-SF proteins across all organisms for each TF-SF harbouring both TFs and nTFs.

Figure B shows the boxplot distribution of the number of organisms harbouring a particular OG. Each point in the distribution represents an OG. The dark pink boxplot shows the distribution for TFs and the blue for nTFS.

The heat map in figure C shows the presence / absence of 34194 OGs across 704 eukaryotic organisms clustered using the clustermap() command in python with “ward” as the method. Presence is considered even if a single OG is present. “grey” marks the presence and “off-white” marks the absence of an OG. Each column represents a particular OG  and each row represents a particular organism belonging to a particular kingdom marked on the left by different colours.

### Figure S3

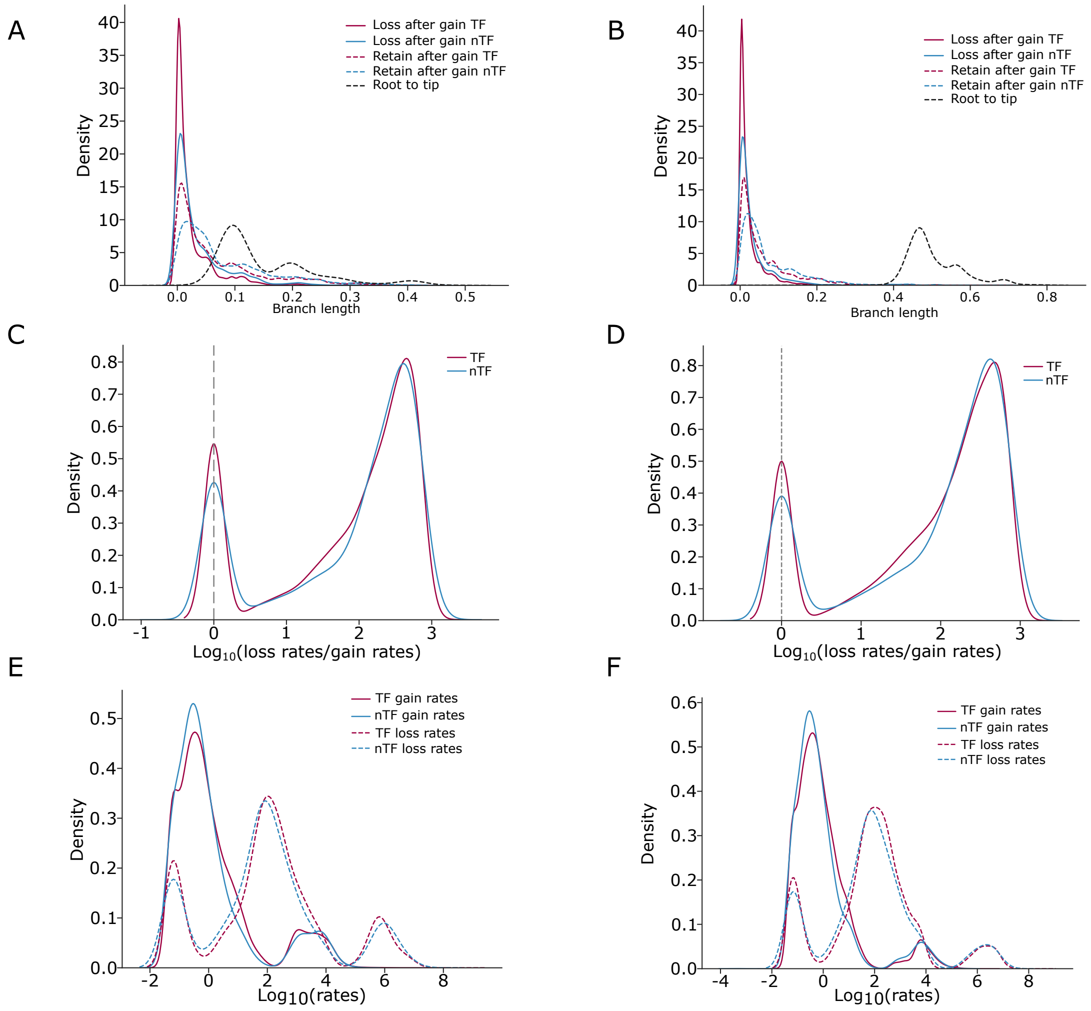


The density plot depicts the branch length distribution for the length travelled by in OG after being gained for the first time for euArch-18S (A) and 18S-archaea (B) trees. The dark pink colour is for TFs and blue is for nTFs. The branch length for loss of OG after its first emergence is denoted by solid lines for both TFs and nTFs. Similarly, branch length for OG being retained after its first emergence is denoted by dashed lines. The black dashed line represents the overall distribution of branch length from root to tip.
Figure C and D represent the distribution of log10 of loss rates/gain rates for each OG for euArch-18S and 18S-archaea trees respectively. The dark pink colour is for TFs and blue is for nTFs. The vertical grey dashed line depicts the value where loss rates and gain rates are equal.

Figure E and F represent the distribution of log10 transition rates for gain (from absence to presence) and (from presence to absence) of OGs for euArch-18S and 18S-archaea trees respectively. The solid lines refer to gain rates for TF (dark pink) and nTFs (blue). The dashed lines refer to loss rates for TF (dark pink) and nTFs (blue).

### Figure S4

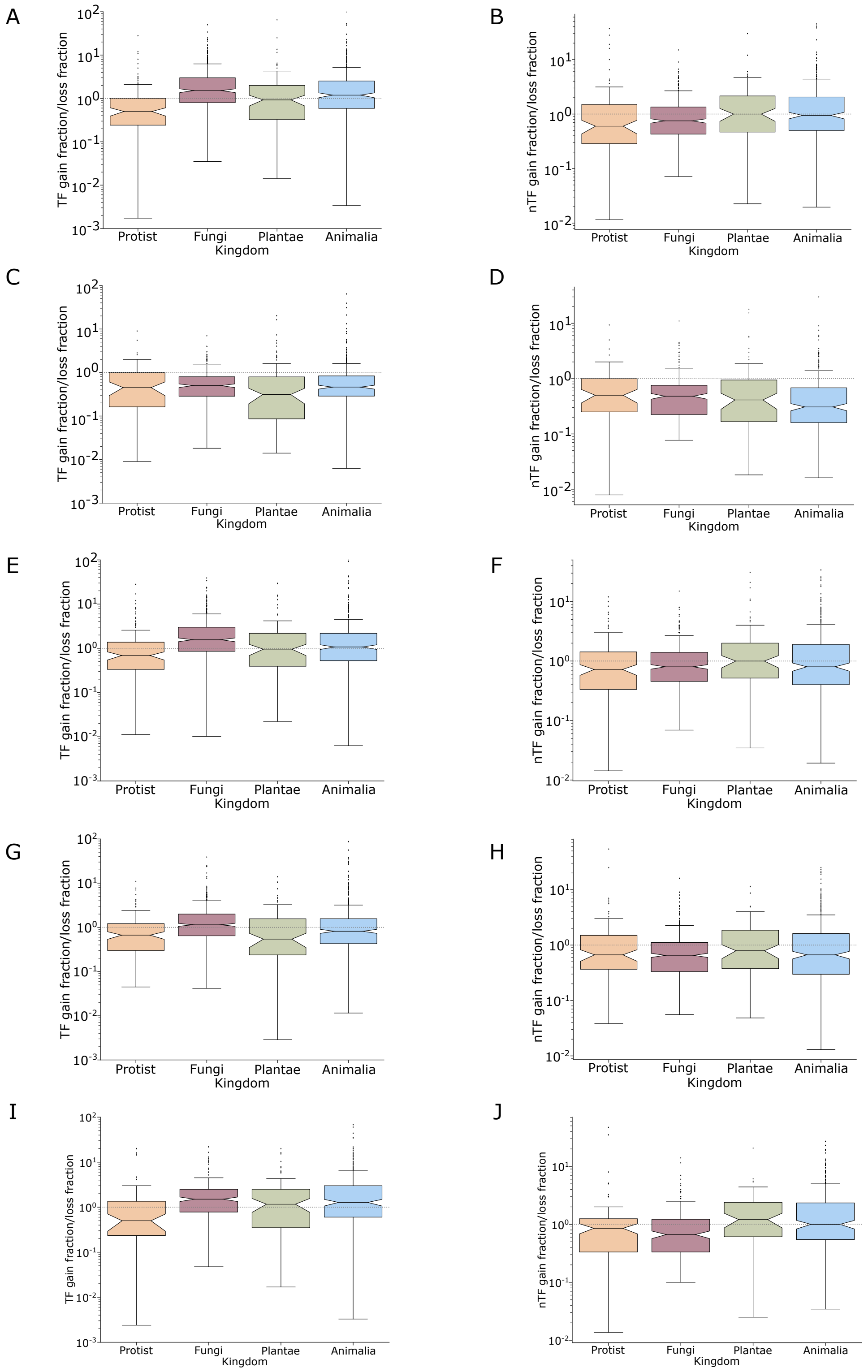


Boxplots represent the ratio of gain fraction/loss fraction (number of OG gained or lost/ total OG in the ancestor node) across the four kingdoms. The figure on the left represent TFs and those on the right represent nTFs for the 4 trees in the following order : euBac-18S (A,B), euBac-concat (C,D), euArch-18S (E,F) and 18S-archaea (G,H). At the kingdom level, for both TFs and nTFs, the gain:loss ratio varies considerably across trees. The euBac-concat tree is dominated by more losses irrespective of the kingdoms or the TF and nTF category (C, D), which is not the case for the 3 18S rRNA trees where there is variation across topologies (A, B, E, F, G, H). Higher loss than gain in the euBac-concat tree does not seem to be the effect of the subset of organisms used in the euBac-concat tree, for the tree with topology similar to euBac-18S tree and for the same set of organisms present in the euBac-concat tree, does not show such a universal dominance of losses (I, J). Thus, some observations of loss:gain rates are not consistent across trees and are affected by its topology and branch length distribution.

### Figure S5

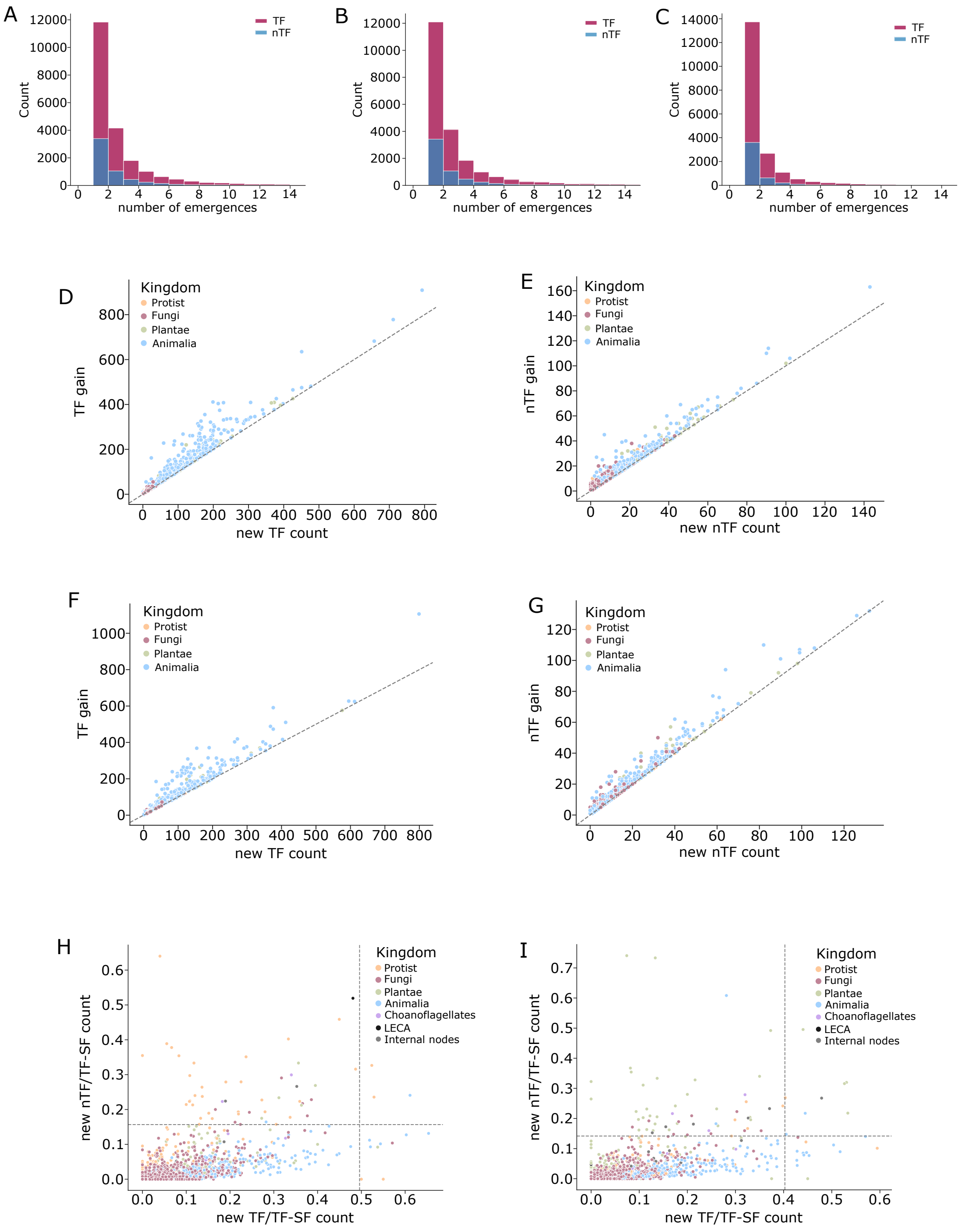


Figure A,B and C represent the histogram for the number of independent emergence for TF (dark pink) and nTF (blue) OGs for the euBac-18S tree, for euArch-18S, 18S-archaea and euBac-concat trees respectively. Here we show a maximum of 14 instances of emergence but it is to be noted that the value goes upto more than 100 emergence for certain OGs.

Scatterplot for gain of new OG versus emergence for new OG at a node was plotted for both TFs and nTFs across the four kingdoms. Figure D and E depict this relation for the euArch-18S tree for TF and nTF respectively.  Figure F and G depict this relation for the 18S-archaea tree for TF and nTF respectively. The dashed grey line marks the 45o line between the x and y axis.

The scatterplots in figure H and I shows the relation between the ratio of new nTF to total TF-SF proteins to the ratio of new TF to total TF-SF proteins for both internal node and tips for the euArch-18S and the18S-archaea tree respectively. The vertical grey line marks the  Q3 + 3 * IQR value (outliers for the distribution) for the x axis (TFs) and the horizontal grey line marks the same for y axis (nTFs).

### Figure S6

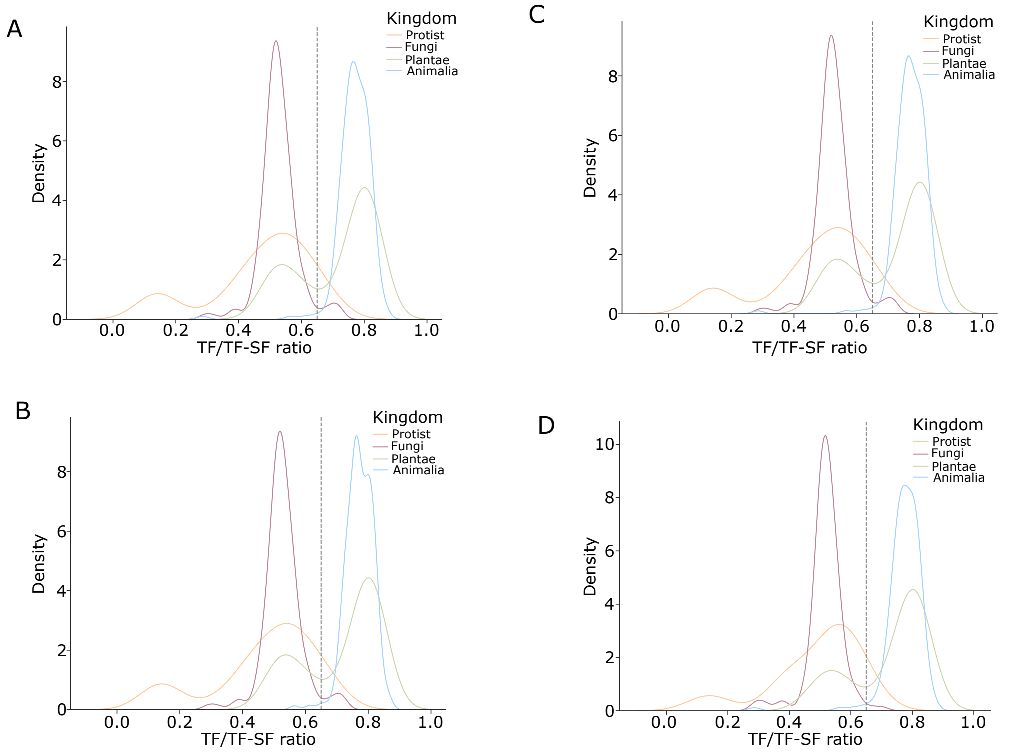


The density plots A,B,C,D shows the distribution of TF to total TF-SF proteins ratio for each extant organism across the four kingdoms marked in different colours,euBac-18S , euArch-18S, 18S-archaea and euBac-concat tree  respectively.

### Figure S7

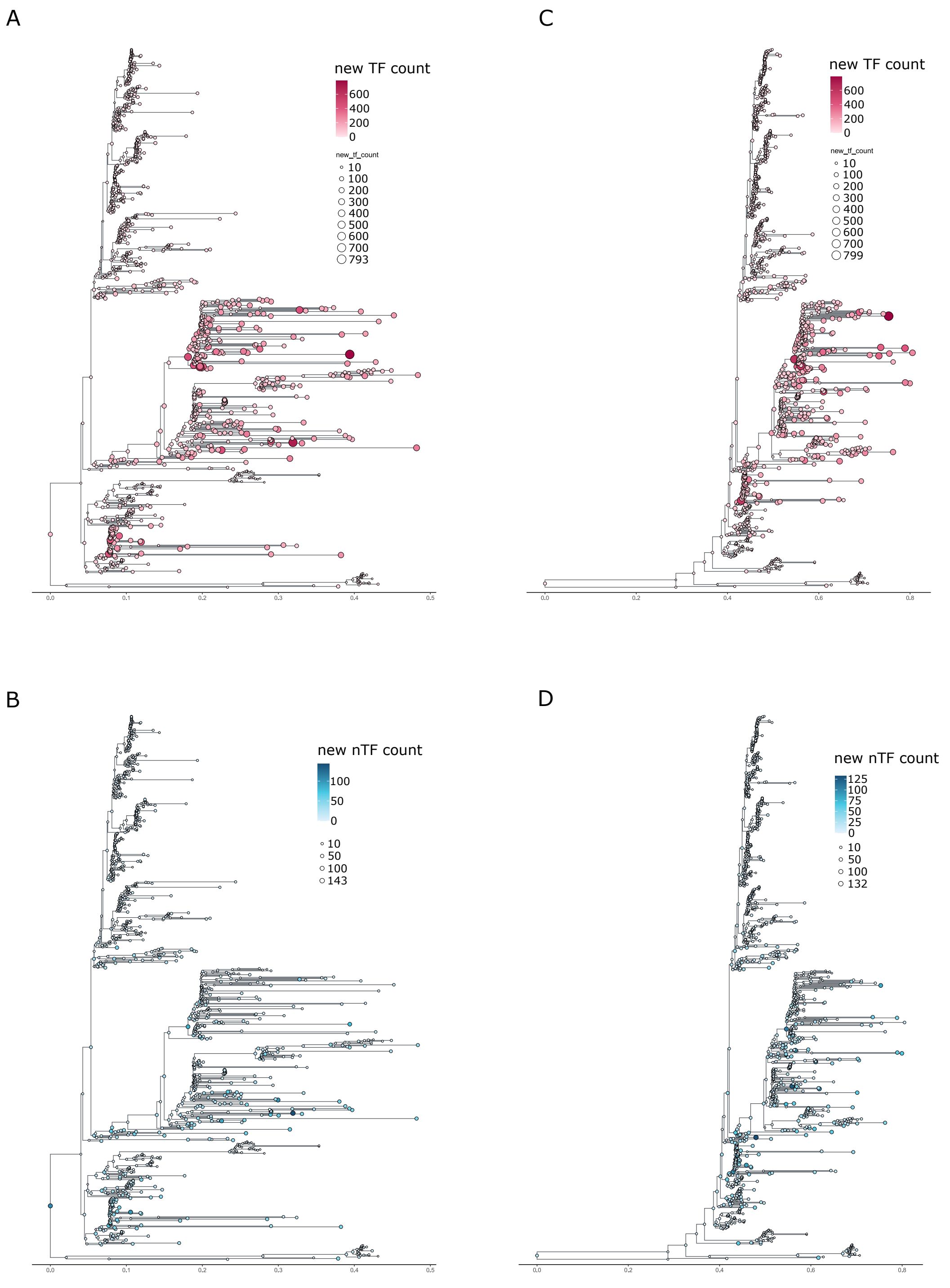


The figure shows the number of innovations (first emergence) of OGs occurring at each node on the phylogenetic tree. Figure A and B depict the number of innovations for the euArch-18S tree for TF and nTF respectively. Figure C and D depict the number of innovations for the 18S-archaea tree for TF and nTF respectively. The dark pink colour represents the TF and blue represents nTF. The darker shades for both TF and nTF represent high values of innovations at a node and lighter shades represent low values.

The same goes for the size for the bubbles with bigger size bubbles exhibiting high values and smaller bubbles showering low values. The topologies for the trees are the same as mentioned in the figure. 3 B and C respectively for the euArch-18S and the 18S-archaea tree.
